# MethPanel: a parallel pipeline and interactive analysis tool for multiplex bisulphite PCR sequencing to assess DNA methylation biomarker panels for disease detection

**DOI:** 10.1101/2020.02.09.941013

**Authors:** Phuc-Loi Luu, Phuc-Thinh Ong, Tran Thai Huu Loc, Dilys Lam, Ruth Pidsley, Clare Stirzaker, Susan J. Clark

## Abstract

**Background:** Multiplex bisulphite PCR sequencing is a convenient and scalable method to comprehensively profile DNA methylation at selected loci. The method is useful for validation of methylation biomarker panels on large clinical cohorts, as it can be applied to DNA isolated from fresh tissue, archival formalin fixed paraffin embedded tissue (FFPET) or circulating cell free DNA in plasma. However, successful clinical implementation of DNA methylation biomarkers for disease detection using multiplex bisulphite PCR sequencing, requires user-friendly sample analysis methods and a diversity of visualisation options, which are not met by current tools.

**Results:** We have developed a computational pipeline with an interactive graphical interface, called *MethPanel*, in order to rapidly analyse multiplex bisulphite PCR sequencing data. *MethPanel* comprises a complete analysis workflow from genomic alignment to DNA methylation calling and supports an unlimited number of PCR amplicons and input samples. Moreover, *MethPanel* offers important and unique features, such as calculation of a polymorphism score and bisulphite PCR bias correction capabilities. *MethPanel* is designed so that the methylation data from all samples can be run in parallel on either a personal computer or a high performance computer. The outputs are also automatically forwarded to a shinyApp for convenient display, visualisation and sharing data with collaborators and clinicians. Importantly the data is centralised in one location, which aids storage management.

**Availability and Implementation:** *MethPanel* is freely available at https://github.com/thinhong/MethPanel

**Conclusion:** *MethPanel* provides a novel parallel pipeline and interactive analysis tool for multiplex bisulphite PCR sequencing to assess DNA methylation marker panels for disease detection.

## Introduction

DNA methylation is the covalent addition of methyl groups to DNA and is associated with essential biological processes. Analysis of DNA methylation patterns allows researchers to explore embryonic development, disease genesis and progression [1]. Since DNA methylation states are highly consistent and stable they are increasingly utilised in clinical diagnostics[2]. Several promising assays to measure DNA methylation are available, including bisulphite PCR sequencing, enrichment bisulphite sequencing, mass spectrometric analysis of DNA methylation, and bisulphite pyrosequencing. A recent community-wide benchmarking study comparing these methods, showed that bisulphite PCR sequencing had the best all-round performance for measuring DNA methylation in low-input clinical samples [3]. Another advantage of bisulphite PCR sequencing is that it can be multiplexed to target multiple regions simultaneously, which is a critical requirement for clinical applications of DNA methylation biomarker panels[4]. Multiplex bisulphite PCR sequencing can also accurately measure methylation in archival, low-input FFPET DNA [5] and may therefore be applicable to low-input circulating cell free DNA (cfDNA) in plasma, a promising noninvasive diagnostic resource [2].

While the laboratory techniques are mature, implementation of epigenetic markers in clinical practice requires the development of user-friendly, large-scale sample analysis and a diversity of visualisation options, which are not met by current tools. For instance, BSPAT [6] and *ampliMethProfiler* [7] can process raw data in FASTA or FASTQ format, but lack an interactive interface for users to modify the format of outputs. *Methpat* can produce interactive plots, but it only focuses on visualisation of DNA methylation patterns and requires BED files that users have to manually produce from the Bismark aligner [8]. *TABSAT* performs all levels of analysis and has an interactive interface, but it is optimised for data generated from Ion Torrent and Illumina MiSeq platforms [9]. *EPIC-TABSAT*, a powerful tool upgraded from *TABSAT* can generate multiple interactive plots and statistics, but due to the nature of its end-to-end web-based service, it is reported to work best with a maximum of 150 targets and 50 samples [10].

Here, we present *MethPanel*, an easy-to-use tool for analysis and visualisation that has been optimised for multiplex bisulphite PCR sequencing data. The tool covers a complete analysis workflow from alignment to DNA methylation calling. *MethPanel* does not limit the number of targets or samples, and offers important but rarely provided features, such as calculation of a polymorphism score and correction for bisulphite PCR bias, that are especially important for the analysis of tumour samples and cfDNA. *MethPanel* can be run on personal computers or high performance computing machines such as Portable Batch System (PBS), Sun Grid Engine (SGE) and Slurm, which allows multiple samples to be run in parallel in a resumable manner. Outputs are automatically forwarded to a shinyApp for visualisation where users can effortlessly customise and share their full data and plots among collaborators easily and privately.

## Results

### Workflow and input data

The workflow of *MethPanel* was built with *Bpipe* [11] in the Linux operating system (Ubuntu 18.04 LTS) as it allows all modules to run automatically, in parallel by samples and to be resumable to generate results in separate directories.

*MethPanel* requires 3 input files: (1) an annotation file of the panel; (2) a plain text file listing all sample names; (3) a config file of trimming and alignment parameters. *MethPanel* supports the human reference genome version hg18, hg19 and hg38.

### Quality control and alignment

In *MethPanel* FASTQ files are trimmed with *TrimGalore* (https://github.com/FelixKrueger/TrimGalore) to produce high quality reads and the percentage of trimmed and remaining reads for each sample are reported. *Bismark* [12] is used in the alignment module with user-defined alignment parameters. To reduce computational memory requirement, *MethPanel* only maps the trimmed reads to the amplicon regions supplied in the user’s annotation file. Internet connection is required to obtain the indexed reference of the genome. A report is generated that describes the number of initial reads inputted, number of mapped reads, number of filtered mapped reads (reads with base quality <30 and length <20bp), depth of CpGs, the number of dropout CpGs (coverage <100x) and amplicon (which has any CpG dropout). Average methylation levels of CpGs and non-CpG cytosines for each sample are also reported. To filter reads that failed bisulphite conversion, users can define the number of non-CpG cytosine residues permitted per read in the config file.

### DNA methylation analysis, pattern calling and PCR bias correction

*MethPanel* performs DNA methylation calling for each sample and merges the results into a complete table of all samples. In this table, each row represents a CpG, each column represents a sample including DNA methylation ratio, the number of methylated reads and coverage. *MethPanel* then performs calling of DNA methylation patterns (rather than just average methylation across an amplicon) and computes a polymorphism score for each sample. For k CpGs in an amplicon, all combinations of 2^k^ patterns are shown in a matrix plot. Polymorphism score is computed as 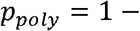 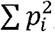, of which p_i_ is the percentage of pattern i [13]. An optional step is to compute PCR-bias values based on the equation 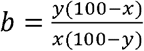 [14], with x corresponding to the expected methylation value and y to the proportion of methylated DNA observed in the ‘methylated-control’ DNA samples. With the assumption the uncorrected curve is hyperbolic curve, DNA methylation is corrected as 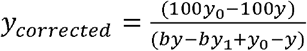, of which *y*_0_ and *y*_1_ are the expected minimum and maximum DNA methylation, respectively [15].

### Visualisation with shinyApp

Table outputs in the previous stages are automatically forwarded to a shinyApp server accompanying the tool. Each user will be given an account with a password to ensure the privacy of their projects. The shinyApp visualisation is summarised as follows. First, the trimmed FASTQ metrics table is displayed and the details of these metrics can be downloaded for further examination. Second, the alignment metrics are shown in the table and can be visualised in the same tab. Third, the DNA methylation tab includes 3 pages, which are (i) an overview page that displays heatmaps for coverage and DNA methylation ratio, (ii) an amplicon DNA methylation page that displays boxplots for both single amplicons and all amplicons and (iii) a DNA methylation of the study group page where users can visualise amplicons by study group for comparison. Fourth, the within-sample pattern of DNA methylation is shown in the heatmaps. Fifth, polymorphism scores are plotted for all amplicons. Finally, *MethPanel* allows users to correct PCR bias by amplicon and CpG site. Corrected data is displayed and analysed in the same format as in the DNA methylation tab.

## Conclusion

We have successfully developed *MethPanel*, a user-friendly tool to analyse multiplex bisulphite PCR sequencing data. *MethPanel* can automatically run multiple samples in parallel and report DNA methylation levels, DNA methylation patterns and polymorphism scores for all amplicons. A variety of interactive plots can be generated with the shinyApp, which users can modify according to their needs without programming knowledge and software installation. The shinyApp server also centralizes the data making for storage management. Moreover, *MethPanel* can handle large numbers of samples and unlimited number of amplicons with interactive high resolution visualisation options. This tool can assist researchers and clinicians to explore DNA methylation data generated from multiplex bisulphite PCR sequencing of clinical samples, including in the analyses of cfDNA.

## Acknowledgements

We thank Jenny Song and Wenjia Qu from the Clark Lab, who performed the multiplex PCR methylation sequencing assays that were used in this paper to demonstrate the utility of *MethPanel*. This work is supported by National Health and Medical Research Council (NHMRC) Fellowship (#1063559) (SJC), Cancer Institute NSW Translational Program Grant # TPG172146 (SJC), Cancer Council NSW RG-18-09 (RP, SJC), National Breast Cancer Foundation IIRS Grant (NBCF-IIRS-18-137) and National Foundation and Medical Research and Innovation (NFMRI) grant (CS) and computational resources provided by the Australian Government through NCI Raijin under the National Computational Merit Allocation Scheme 2019, project wk73 (SJC, PLL).

---

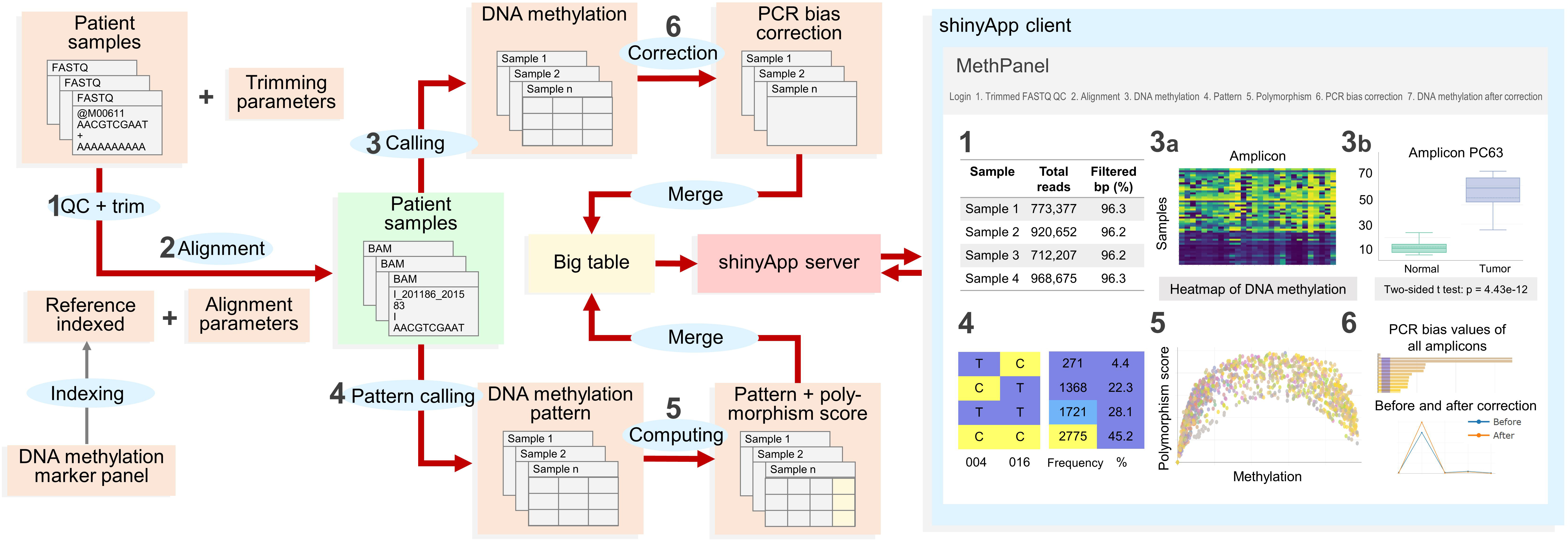

